# High resolution *in situ* structural determination of heterogeneous specimen

**DOI:** 10.1101/231605

**Authors:** Benjamin A. Himes, Peijun Zhang

## Abstract

Macromolecular complexes are intrinsically flexible and often challenging to purify for structure determination by single particle cryoEM. Such complexes may be studied *in situ* using cryo-electron tomography combined with sub-tomogram alignment and classification, which in exceptional cases reaches sub-nanometer resolution, yielding insight into structure-function relationships. All maps currently deposited in the EMDB with resolution < 9 Å are from macromolecules that form ordered structural arrays, like viral capsids, which greatly simplifies structural determination. Extending this approach to more common specimens that exhibit conformational or compositional heterogeneity, and may be available in limited numbers, remains challenging. We developed **emClarity**, a GPU-accelerated image processing package, specifically to address fundamental hurdles to this aim, and demonstrate significant improvements in the resolution of maps compared to those generated using current state-of-the-art software. Furthermore, we devise a novel approach to sub-tomogram classification that reveals functional states not previously observed with the same data.

The software is freely available from https://www.github.com/bHimes/emClarity

Tutorial documentation and videos at https://www.github.com/bHimes/emClarity/wiki

## Introduction

Recent advances in the capabilities of cryo-electron microscopy (cryoEM) are changing how we think about structural determination of complex biological assemblies. In addition to reaching atomic-resolution, the development of advanced classification techniques, such as maximum-likelihood classification as implemented in FREALIGN^1^ or RELION^2^, has enabled possibilities for probing macromolecular functional dynamics using a single particle analysis (SPA) approach. For a sample to be suitable for SPA, it must yield tens to hundreds of thousands^3^ of individual instances of biological macromolecules, commonly referred to as particles, which must first be purified to relatively high compositional and conformational homogeneity^4^ and subsequently imaged in many different orientations. These two conditions are often difficult to achieve, especially for the large assemblies of biological complexes represent the functional context most relevant to cellular activities^5^. When this is the case, cryo-electron tomography (cryoET) is the preferred approach, capable of generating three-dimensional (3D) reconstructions of pleomorphic samples *in situ*.

These reconstructions, called tomograms, are generally limited to 3-4 nm resolution. This limit on the resolution of a tomogram is largely due to the extremely limited electron dose chosen to prevent excessive radiation damage to samples during the collection of many projection images. Additionally, the signal that is available in these noisy images, is not distributed evenly in the tomogram, resulting in anisotropic resolution, a consequence of both increasing specimen thickness at high tilt angles and the mechanical limits imposed by the objective lens, which restricts the angular sampling to +/− 60°. The resulting distortion is colloquially referred to as the “missing-wedge effect” named for the shape of the sub-sampled region in Fourier space of a single-tilt-axis tomogram.

These resolution-limiting issues can be overcome when many copies of a macromolecule are present in a tomogram, extracted *in silico*, aligned iteratively, and averaged using procedures that share many similarities to SPA, *viz*. cryo-electron sub-tomogram alignment and classification (cryoSTAC). Such averaging can increase the signal to noise ratio (SNR) in the final map and “fill-in” the missing wedge by completing the angular sampling, provided that missing-wedge-introduced bias during image processing can be mitigated. This is generally accomplished by explicitly considering the contribution of each sub-tomogram to the final average as a function of spatial frequency in Fourier space, by modifying the (dis)similarity metric used for alignment and/or classification; the two most common being the constrained cross correlation^6,7,8^ and the Euclidean distance^9,10^. The extent of information transfer during this process is described by 3D Contrast Transfer Function (3D-CTF). The 3D-CTF can describe sampling that is affected by total radiation exposure, ice-thickness as a function of tilt angle, weighting in the tomogram reconstruction process, or a quality factor that reflects the accuracy in orientation determination and the angular sampling permitted by the microscope. The reader is referred to recent reviews which cover both the general principles^11^ and computational approaches^12^ in greater depth.

Obtaining averages of sub-tomograms at low resolutions, 15-20 Å, is routine and necessary for further characterization of the data via statistical methods, which then describe the population based on physical differences rather than spatial (orientational) differences. This process, commonly referred to as “classification”, allows for separation of multiple biological states from the same sample allowing for *in situ* structural determination of functional conformations^13,14^. Compared to SPA, this is arguably the greatest strength of cryoSTAC, because each particle exists as a unique, albeit distorted, 3D reconstruction. This allows for a direct analysis of the 3D variance, the value of which has been discussed extensively^15,16,17^. To date, only a few structures have been solved at resolutions better than 10 Â using cryoSTAC^18,19,20,10,21,22,23^, a critical threshold beyond which flexible molecular fitting approaches are more reliable^24^. This approach of integrating high resolution data into medium resolution maps has proven to be very useful in understanding dynamic and often transient complexes ^19,13,25^ such that reaching this resolution is of great interest.

We present here a complete set of GPU-accelerated programs called **emClarity** for **<u>e</u>**nhanced **<u>m</u>**acromolecular **<u>cl</u>**assification and **<u>a</u>**lignment for high-**<u>r</u>**esolution ***<u>i</u>****n situ* **<u>t</u>**omograph**<u>y</u>**, with the aim of reaching beyond the critical sub-nanometer resolution in cryoSTAC for diverse specimens. We have focused our efforts on those areas of image processing that are likely to yield the greatest improvements, as suggested by empirical observation and theoretical calculations^26,20,27,28^: accuracy of tilt-series alignment, improved defocus determination and CTF correction, explicit treatment of anisotropic resolution, and more robust classification.

## emClarity guiding principles and workflow

emClarity has been built with a strong emphasis on reliability, taking several measures to minimize the fitting of noise that may be introduced through iterative alignment procedures^29^. In particular, emClarity uses a robust method of alignment (the “gold standard” approach^30^), significantly reducing fitting noise at minor computational cost^31^. Specifically, during alignment, the data are weighted according to the spectral signal-to-noise ratio (SSNR) as determined using the “gold-standard” Fourier Shell Correlation (FSC) based on a modified version of the figure-of-merit (FOM)^32^ which draws on ideas from x-ray crystallography. In emClarity, the FOM limits information beyond resolutions beyond where the 1-bit information curve intersects the FSC^33^. A typical cryoSTAC workflow is illustrated in figure 1, with the most impactful of our enhancements highlighted in red text. The addition of a branch in the iterative process to refine the tilt-series alignment is not available in other image processing packages.

**Figure 1.**
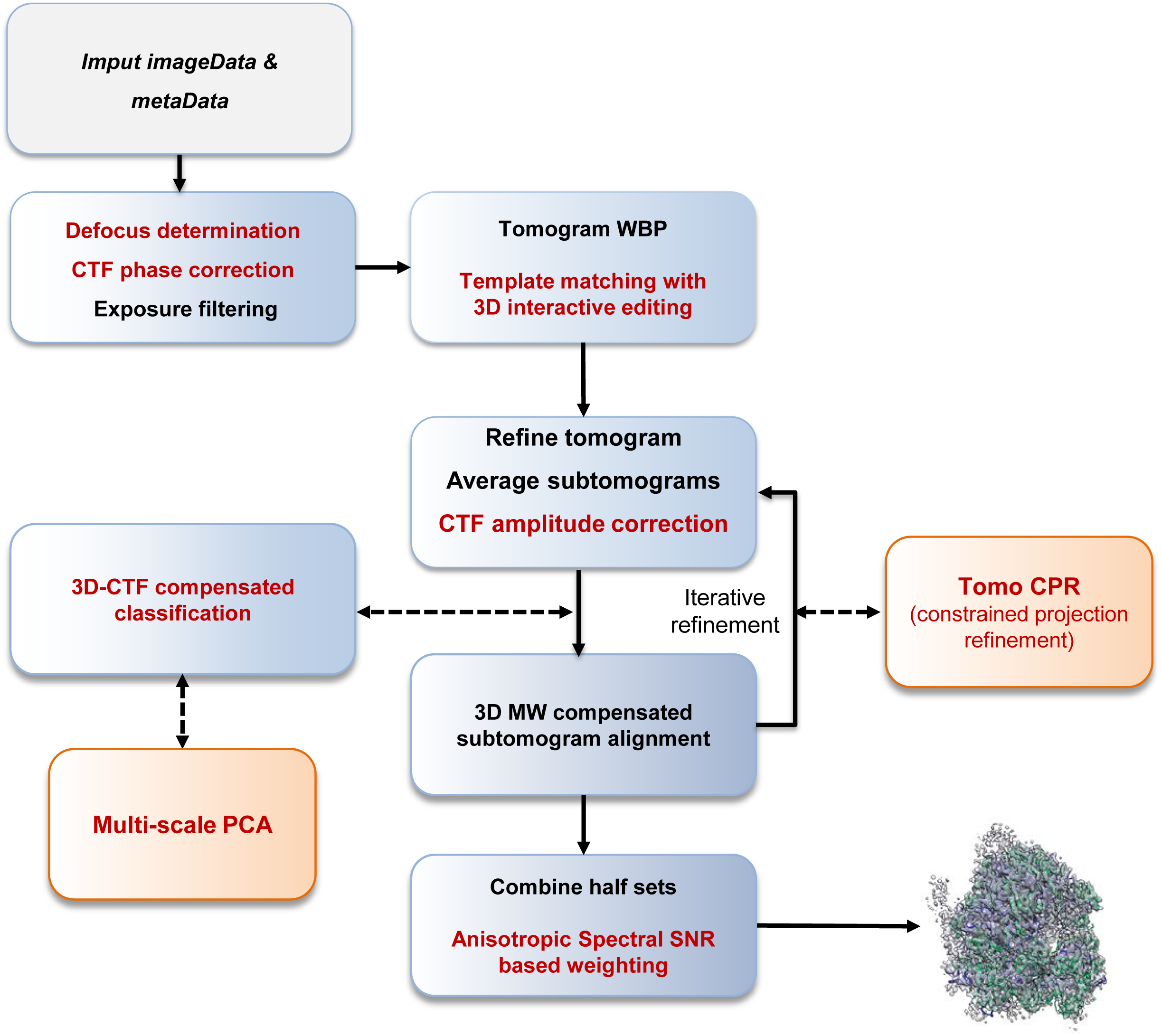
The emClarity workflow for cryoSTAC, where significant improvements introduced in emClarity are highlighted in red text, while entirely novel additions to the process are indicated in orange boxes.

## Major features for improving resolution in emClarity

In each of the following sections, we describe the principles underlying each of the major features in emClarity, with algorithmic and mathematical details left to the supplemental methods.

### Refinement of tilt-series alignment

A prerequisite for 3D reconstruction of a tomogram is the refinement of the projection geometry, including tilt-axis angle, in-plane shifts and rotations, magnification, tilt angle, and possibly other distortions like non-perpendicularity of the electron beam or a skew between the x & y axes^34^. This process (tilt-series alignment) is most commonly accomplished by using gold beads as high-contrast fiducial markers. Additional approaches, based on locating and tracking image features^35,36,37^ or on projection matching using an intermediate tomogram as a 3D model^8^ are available, but their success is generally sample-dependent^11^ and require significant user input, which has been somewhat ameliorated by automation with the recently released Appion package^38^.

We have integrated into emClarity a novel algorithm called tomo-CPR (tomogram constrained particle refinement) for the iterative refinement of the tilt-series alignment using an approach that shares some similarity with the “particle polishing^39^” implemented for SPA in RELION with two primary differences. First, in generating reference projections for orientation determination, tomo-CPR includes information from neighboring particles as well as non-particle information that is relevant for localization as illustrated in supplemental figure 1. Second, tomo-CPR constrains neighboring particles to behave similarly within a given projection as well as requiring them to vary smoothly as a group from projection to projection through the tilt-series.

### Maximizing weak signal in the reconstructions

Real space masks that follow the particle envelope are useful in the maximization of the SSNR, however, care must be taken to avoid inadvertently introducing correlations by application of the same real space masks to data from each half-set^42^. We apply a simple but effective iterative dilation starting from the highest SNR pixels to those at the mean while enforcing connectedness. This means that flexible regions of macromolecules, or complexes that have variable occupancy with time are retained by using such a masking approach that is based on connectivity.

In reciprocal space, image features are grouped by spatial frequency, such that masks can be applied to specific resolution bands. High resolution features degrade at a faster rate as a function of electron exposure, and so we have adapted for projections of tilted specimens the optimal exposure filter^41^ originally described for SPA. Briefly, the exposure-based filter weights each projection according to the cumulative electron dose, with the goal of maximizing the SNR in the final weighted sum. To avoid any low-pass filtering effects, the final sum is re-weighted according to how much information has been included as a function of spatial frequency. While we might consider applying this to the projections directly, as with CTF amplitude restoration, it is safer to reweight the final reconstruction (sub-tomogram average) to avoid inadvertently amplifying noise.

### Improved defocus determination for more accurate CTF correction

Even with the advent of direct electron detectors, estimating the defocus in an image with a dose of only 1-2 e^-^/Å^2^ requires meticulous microscope alignment, optimized data collection schemes and a sample that provides abundant signal^18^. For cases where these conditions are not met, we have devised a new algorithm to maximize the information available for estimating the defocus value. For the initial defocus determination, we apply rotational averaging to the power spectrum as estimated from periodogram averaging, as described previously^43^. After obtaining this primary estimate of the mean defocus at the height of the tilt-axis, we then fit a 2D astigmatic function to the average power spectrum without rotational averaging, as is routine in SPA^44^ using a low pass version of the power spectrum for background subtraction^45^.

The number of periodograms available for coherent averaging decreases in tilted images as a defocus gradient is introduced perpendicular to the tilt-axis. To increase the number of periodograms available, the known defocus gradient may be used to resample the portion of the power spectrum between the first two zeros of the CTF^46^, an approach currently used in IMOD to enhance the accuracy of the global defocus determination. We devised a novel approach which extends this concept to resample *all* the CTF information by using an approximate form of the phase aberration equation and taking advantage of properties of the discrete Fourier transform as justified in the supplemental methods.

### Statistical optimization of the SNR in the final map

Several programs for correcting the CTF phase and amplitude modulations directly in the 2D projections of tilted images are available; the two predominant being CTFPLOTTER and CTFPHASEFLIP^46^ included in the IMOD package^47^, and TOMOCTF^43^. They differ mainly in their approach to background subtraction and amplitude restoration, the former relying on an approximate correction based on the detectors MTF, and the latter implementing a novel filter that acts similarly to a Wiener filter while tuning the strength of signal dampening in three different regions of frequency space. Both approaches are sub-optimal as they rely on restoring amplitudes in the Fourier transforms of individual projections which are particularly noisy, and thereby reduce the fidelity with which the CTF amplitude modulations may be restored^48^.

A more attractive approach is to correct the phases on the projections, and then to address the amplitudes after building the 3D reconstruction (sub-tomogram average). This is the approach taken in SPA and also in the adaptation of RELION for sub-tomogram averaging^10^ which uses a Wiener-like filter to restore the amplitudes based on an estimate of the SSNR(q) as an integral part of the underlying statistical model. A typical Wiener filter based approach has also been described recently in the near atomic resolution structure of the HIV capsid protein^18^ but is only implemented through “in-house” software.

One potentially limiting factor in the RELION implementation is the reliance on the 3D-CTF model for the phase correction of the individual particles. This means that the 3D-CTF must be calculated (and stored on disk) for every particle, which we suggest is a poor choice for imaging with a defocus range of 2-4 μm as is common in cryoSTAC. While disk storage is relatively cheap, a 3D-CTF of ~512^3 pixels is needed for every particle to prevent significant aliasing of the CTF^49^. We opt to correct for the phase inversion on the projection images, oversampling the tiles to avoid any aliasing, and then constructing a full 3D-CTF model which is used in amplitude restoration of the reconstructed density. Our 3D-CTF also takes into consideration the R-weighting that is applied during tomogram reconstruction. Considering these two differences, we then calculate a set number, usually nine, 3D-CTFs evenly distributed perpendicular to the tilt-axis for each tomogram which are then incorporated into the final re-weighting by an adaption of the “volume normalized single-particle Wiener Filter^50^”. Importantly, this adaptation involves explicitly accounting for directional anisotropy in the distribution of signal.

### 3D-CTF compensated classification

We use multi-variate statistical analysis (MSA) for classification. This involves first defining a set of descriptors or *features* which are then searched for patterns that are common to a sub-group called a pattern class, or simply a *class*. Grouping inputs by these patterns is accomplished with a clustering algorithm; commonly k-means, hierarchical ascendant classification (HAC) or a neural networks approach via a self-organizing map (nn-SOM), all three are implemented in emClarity. Since the “missing-wedge” produces significant artifacts that are specific to the orientation of each particle in the sample, but not necessarily its identity or conformation, it is challenging to resolve meaningful patterns in cryoSTAC data. Estimating the effect of the missing-wedge by using a binary mask has been shown to be a good first order correction called wedge masked differences (WMDs)^54^. We replace this mask with our 3D-CTF function which results in a more accurate estimate of the artifacts introduced by the “missing-wedge” and allows for higher resolution information to be considered in the clustering.

### Multi-scale clustering

In a naïve approach, a clustering algorithm will interpret every individual voxel as an independent measurement along one dimension of an N-dimensional space, where N is the number of voxels in each sub-tomogram at the time of analysis. To reduce the impact of the “missing-wedge” and to introduce a correlation between pixels, a smoothing filter is typically applied to the data prior to clustering. In effect, this tells the clustering algorithm that neighboring pixels are measurements of related features in the full sample space. Because our 3D-CTF WMD feature vectors are robust to the “missing-wedge” at higher resolutions, we extend this idea to encode *a priori* biological information by forming them from data that have been convolved with Gaussian kernels which introduce inter-voxel correlations at biologically relevant length scales. For example, ~ 10 Å for alpha-helical density, 18-20 Å for RNA helices or small protein domains, and ~ 40 Å for larger protein domains. This approach is similar to existing ideas in multi-scale multi-variate statistical analysis, that use the discrete wavelet transform with a limited set of coefficients followed by clustering analysis of the data reconstructed independently using a limited subset of wavelet basis^55^. In our case, each Gaussian kernel can be viewed as a simple wavelet localized at the origin, and defined in frequency by the biological length-scales previously mentioned. The primary difference is that the coefficients obtained by projection of the data along the direction of the largest singular vectors at each of these lengths scales are then concatenated into a single matrix, and optionally weighted relative to each other, such that the clustering algorithm, whether it is k-means, HAC, or nn-SOMs, ***considers each length scale simultaneously***, providing a much richer description of the feature space. In effect, this teaches the clustering algorithm to “see the forest for the trees.”

## Results

### emClarity improves resolution in sub-tomogram averaging

Given the inherent difficulty in working with extremely low SNR cryoEM data, and the sensitivity of the results to optimal selection of parameters^1^, we have elected to test and demonstrate our software using two publicly available data sets from the Electron Microscopy Pilot Image Archive^56^ (EMPIAR). We show these published/deposited maps, juxtaposed with the maps obtained with emClarity in figure 2. A total improvement in the yeast 80s ribosome from EMPIAR-10045 using RELION version 1.4 (EMD-3228^57^) from 12.9 Å to 7.8 Å is achieved using emClarity (Figure 2a). For the mammalian 80s ribosome from EMPIAR-10064 using pyTOM (EMD-3420^21^), we obtained an improvement from 11.2 Å to 8.6 Å (Figure. 2b).

**Figure 2.**
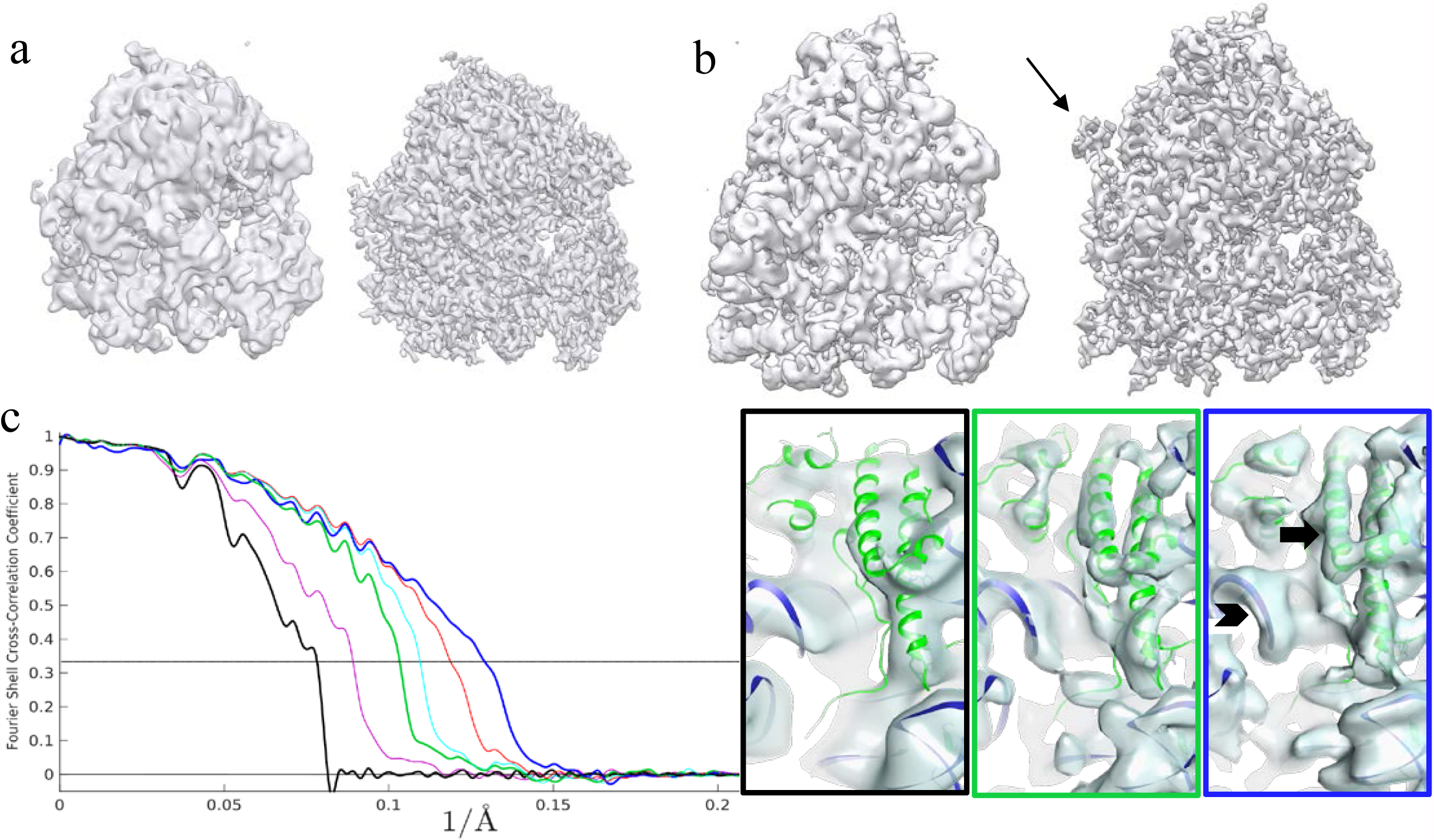
Improvement in resolution of sub-tomogram averaging using emClarity. (a) Comparison of the sub-tomogram average of yeast 80s ribosome by RELION (EMD-3228) (left, at 12.9Å^-1^ resolution) and by emClarity (right, at 7.8Å^-1^ resolution), using the same raw data from the electron microscopy public image archive (3,233 Yeast 80s ribosomes, EMPIAR-10045). (b) Comparison of sub-tomogram averages of rabbit 80s ribosome by pyTOM (EMD-3420) (left, at 11.2Å^-1^ resolution) and emClarity (right, at 8.6Å^-1^ resolution), using the same raw data (1,400 Rabbit 80s ribosomes, EMPIAR-10064). Arrow points to an additional feature present just outside the peptide exit tunnel which is only revealed with the more conservative masking procedure in emClarity. (c) cross-FSC between the sub-tomogram averages by emClarity and the SPR cryoEM map (EMD-2275) of yeast 80s ribosome. The first five curves use orientation parameters from the Relion 1.4 alignment, cumulatively including additional features from emClarity: magenta, improved CTF estimation and correction with the optimal exposure filter; green, adding one round of tomo-CPR; cyan, adding in the per-tilt defocus estimation; red, adding in explicit consideration of resolution anisotropy in the adapted single particle wiener filter. The final dark blue curve incorporates all these features plus the alignment parameters determined from scratch in emClarity. Representative views of sub-tomogram averages, with a rigid body docking of yeast 80s atomic model (PDB-47VR) for visualization. Panel frame colors, black, green and blue, correspond to the cross-FSC plots respectively. The arrow and chevron highlight the resolved alpha helices and RNA structures, respectively.

To evaluate the relative impact of each of the individual features implemented in emClarity, we incrementally included them into several reconstructions of the yeast 80s ribosome. To control for errors in alignment and to have a one-to-one comparison with EMD-3228, we used precisely the same particles and orientation parameters from the star files that accompany the raw data EMPIAR-10045. We compare each map to an external reference map derived from SPA (EMD-2275^58^), via a cross-Fourier Shell Correlation (cross-FSC), starting from the RELION reconstruction as a control (Figure 2c). The results demonstrate the recovery of additional signal from the same data as each subsequent feature is incorporated.

The accuracy of our combined CTF correction approach, phases on oversampled 2D-tiles combined with optimal-exposure filtering and 3D-CTF based Wiener filtering is reflected in the magenta curve in figure 2c, which shows a significant improvement over the cross-FSC of the control, even though they are reconstructions using the same particles and orientations. The largest single improvement comes from the tomo-CPR which is shown in green (and obviously includes the features in the magenta curve as well.) A more modest improvement is measured when we add in a per-tilt defocus estimation using our novel approach to resample periodograms from tilted images, as reflected in the cyan cross-FSC. When we also explicitly consider anisotropy in the SSNR in our adaptation of the single particle Wiener filter, we see another substantial improvement in the cross-FSC in the red curve. The yeast 80s sample that was used has a strong preferential orientation which is reflected in the FSC-cones and a plot of the angular distribution in in supplemental figure 2. The final and highest resolution curve represents an alignment carried out in emClarity with all features added, illustrating the additional impact these advances have on the accuracy in the orientation determination.

In addition to improved resolution, as noted in figure 2b, there is a density outside the peptide exit tunnel of the ribosome (white arrow) that is present in the map derived with emClarity, but not in the map derived with pyTOM. Finally, in figure 2d, we show the density from a peripheral region with a rigidly docked model of the yeast 80s ribosome (PDB-47VR) that underscores the difference in interpretability between the maps derived from the current state-of-the art and emClarity. There is a clear improvement in both RNA and protein structures.

### emClarity improves classification and reveals multiple ribosome functional states

Using multi-scale clustering combined with 3D-CTF compensated Principal Component Analysis (PCA), emClarity reveals subtle conformational differences and distinguishes minor populations from noisy and distorted images, as demonstrated with yeast 80s ribosome data from EMPIAR-10045, and mammalian 80s ribosome data from EMPIAR-10064. Such results were not previously obtainable using existing software^21,57^.

#### Classification of non-translating Yeast 80s ribosomes

The ribosome is a complex molecular machine composed of RNA and protein which exists in many functional states and interacts with an array of co-factors. The major domains are named by their sedimentation coefficients (s, svedberg) where the eukaryotic ribosome is composed of two major domains dubbed the large subunit (60s) and small subunit (40s). While the ribosome has a well-conserved catalytic core which mediates the peptidyl transferase reaction^59^, it is increasingly subject to more complex regulation in higher organisms resulting in an expanded set of both RNA and protein components. RNA expansion segments are found primarily at the periphery of the ribosome and are typically highly dynamic and difficult to resolve in structural analysis. One particularly good example is es27, an approximately 150 ÅRNA helix which predominantly adopts one of two conformations separated by about 90☐, shown in orange in figure 3 a-e. The first situates the end of the RNA helix just outside the peptide exit tunnel on the 60s subunit (es27_pet_, figure 3 a, b, d, e) and the second points toward the tRNA exit site (es27_L1_, figure 3c). This dynamic domain is generally observed in cryoEM maps as a superposition of these two states, as is the case with the currently published results by ML classification in RELION^57^. A notable exception being ribosomes with accessory complexes bound at the peptide exit tunnel, e.g. Sec61, are known to bias the conformation to the es27_L1_^60^.

**Figure 3.**
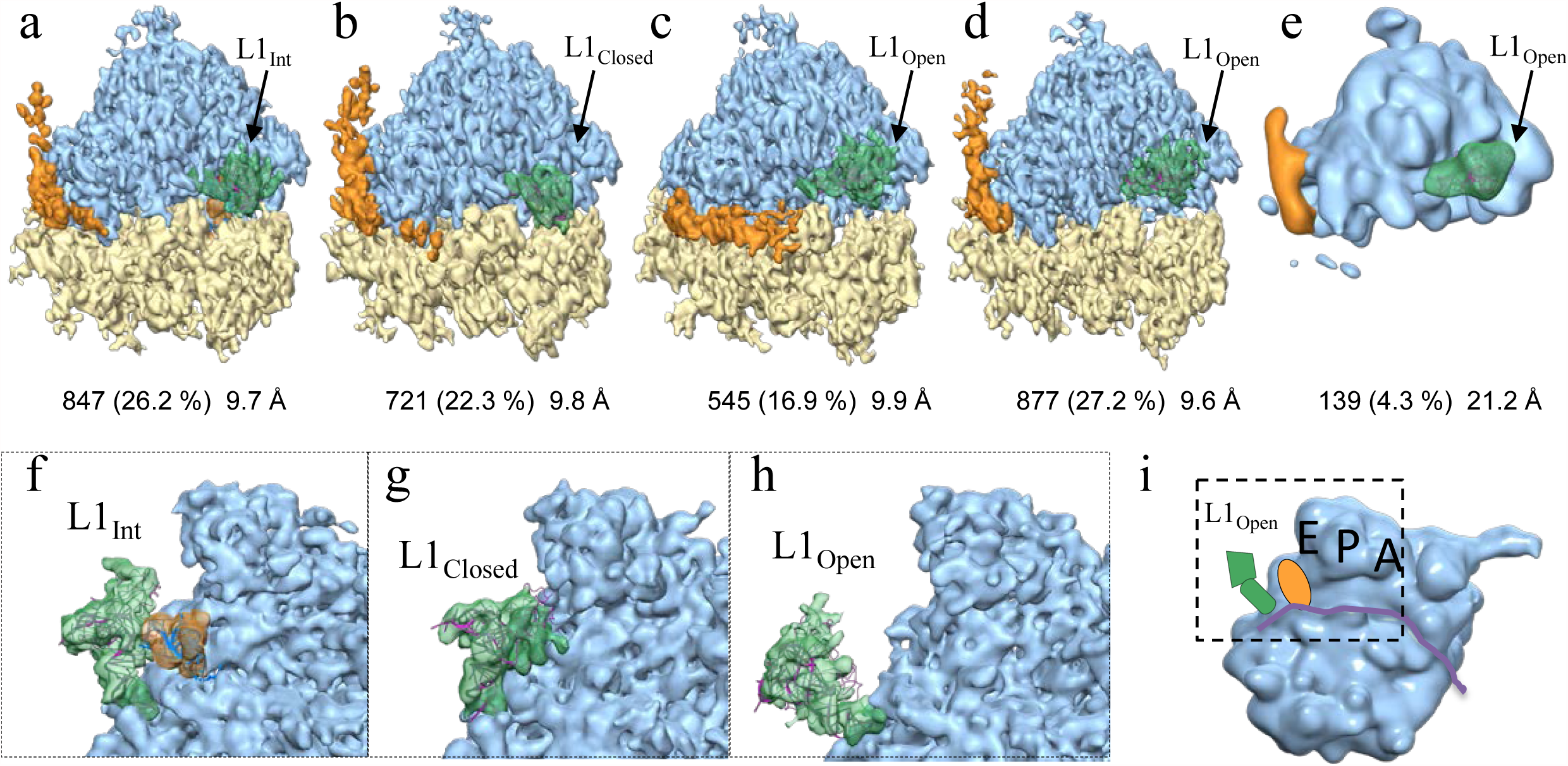
Classification of yeast 80s ribosome (EMPIAR-10045) with 3D-CTF compensated missing-wedge and multi-scale PCA in emClarity. (a-e) Four major classes and a minor class contributing 96.9% of subtomograms are shown with number and percentage of contributing units and resolution indicated below. The remaining 3.1% comprised a 6^th^ minor class with no significant structural features and were removed from analysis. The highly dynamic L1 protuberance (green) and RNA expansion-segment 27 (orange) are captured in distinct conformations in these classes. (i) cartoon illustrating the region displayed in (f-h) from the inter-subunit space with the E,P,A sites labelled and L1 (green) E-site tRNA (orange) and mRNA channel (purple ribbon.) (f-h) Enlarged views with the small-subunit removed for clarity, looking out from the inter-subunit space, of the L1 protuberance in an intermediate position bound to P/E tRNA shown in orange (f, class a), fully closed interacting with the 60s central protuberance (g, class b), and fully open (h, classes c-e) respectively. The riboprotein and selected RNA helix components of the L1 protuberance (rpL1,h76,h79 from PDB 3J78) shown in magenta ribbon after rigid body docking into the respective density.

Another example of a highly dynamic ribosome domain is the L1 stalk – comprised of protein L1, and RNA helices h75, h76 and h79 from the 25s portion of the 60s subunit^61^ – which moves through ~ 55 Å during a translocation cycle. The motions of L1 are well correlated with several defined functional translocational states as observed using single molecule FRET and SPA^62^. Using emClarity, we can discern three of these L1 conformational states isolated from thermal (stochastic) fluctuations of the non-translating yeast 80s ribosome: L1_open_, L1_int_, and L1_closed_ shown in green with variable occupancy in the five classes in figure 3. In addition to isolating dynamic states, identifying very sparsely populated classes is a particularly important and challenging task for classification of cryoEM data. We see in figure 3e the dissociated 60s subunit occupying a minor class, only ~4% of the data set or roughly ~140 sub-tomograms. In contrast, the Maximum likelihood approach implemented in RELION found three classes, one designated as a junk class and two relatively indistinguishable classes^57^. This minor population could only be isolated in the case where feature vectors built from the projection on the principal components from at least three length-scales were simultaneously clustered.

#### Mammalian 80s ribosome

In contrast to the non-translating yeast specimen, the mammalian ribosomes imaged in EMPIAR-10064 were prepared from clarified rabbit reticulocyte lysate using a buffer low in Mg^2+^ but lacking polyamines, such that cofactors should co-purify excepting perhaps some loss of e-Site tRNA^63^. We extracted 3,090 ribosomes from the four tilt-series deposited as the “mixed-CTEM” data set on EMPIAR. emClarity identified five predominant classes as shown in figure 4a-e. Three of these classes show ribosomes adopting a non-rotated 40s conformation with variable tRNA eeF1A occupancy (class I-III), while two very similar classes adopted a mid-rotated (-5-6☐) 40s conformation with eeF2 present (class IV-V).

**Figure 4.**
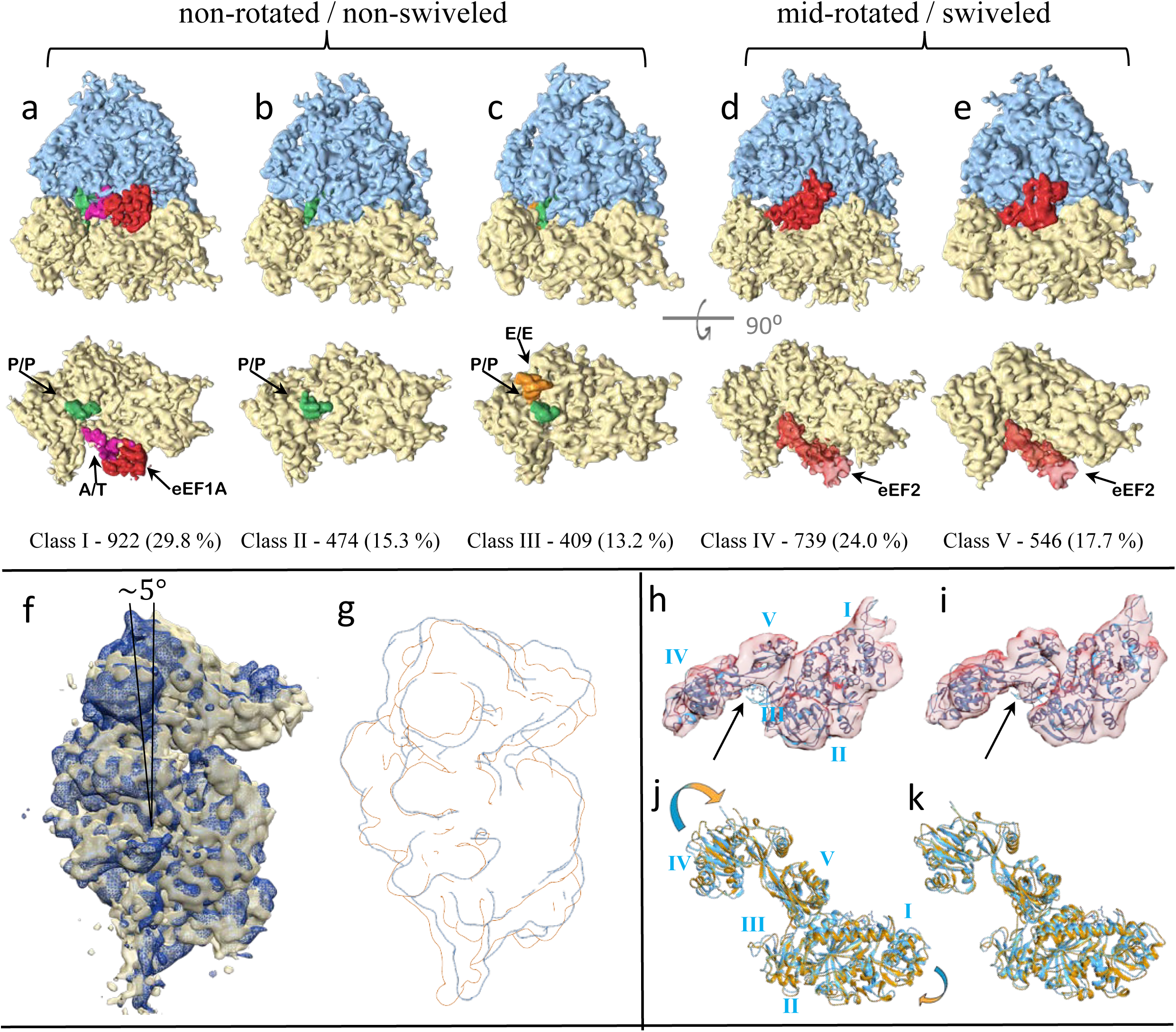
Classification using multi-scale Clustering with 3D-CTF compensated feature vectors reveals five distinct translocational species from 3,100 particles. (a-c) Classes I-III represent a post translocational state with the co-factors shown in the lower row from the inter-subunit surface with the 60s subunit removed for clarity. (d-e) Classes IV-V have a mid-rotated 40s and a swiveled head corresponding to a pre-translocational intermediate. (f) Class IV in dark blue overlaid with class III in gold, showing the midrotated 40s state. (g) Outline from a low-pass filtered overlay as in (f). (h) MDFF of eeF2 (orange) with the density from class-IV starting from PDB-4ujo (cyan ribbon) shows similar conformation in eeF2 domains II, III and V, while eeF2 domain IV deviates the most. (i) Same as (h), but with class V showing smaller deviations. (j-k) The rigid body docking of PDB-4ujo into the density from class IV/V respectively shows overall close agreement, except the stronger density between eeF2 domains III & V in (k). Arrows point to this density which is occupied by the antibiotic Sordarin in PDB-4ujo, but is not present in the sample used in this study.

**Figure 5.**
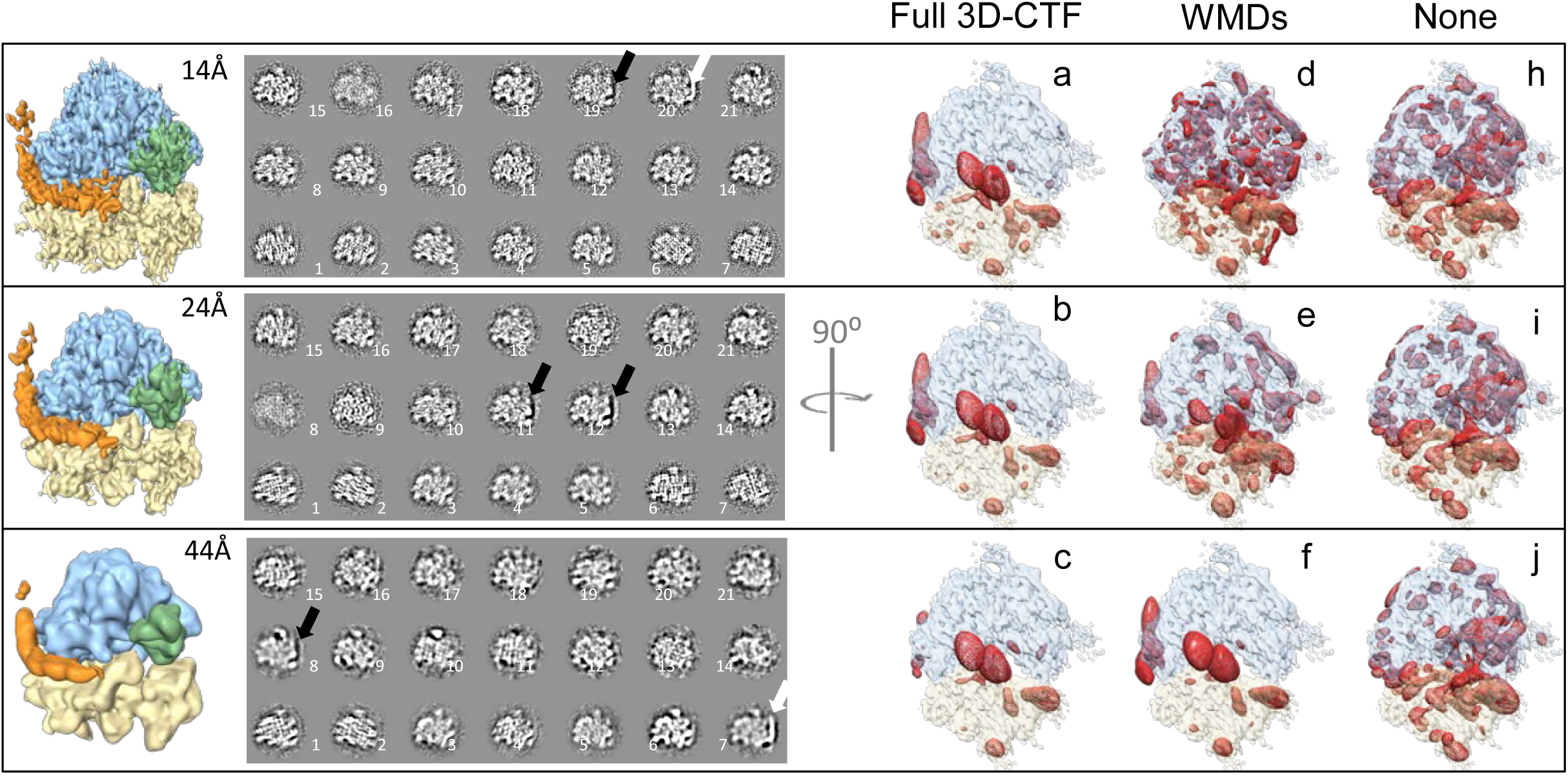
Illustration for the impact and compensation of the missing wedge at multiple length scales. Far left column, the total average filtered by Gaussian kernels of variable width to correlate voxels over the given length scales. Center column, eigen-images composed of the average plus the eigenvector, sorted from the most variance explained (1) to least (21). Black and white arrows show example where es27 density is absent or present in the L1 position. Right three columns show 3D-variance maps in red overlaid with the average as a visual guide. (a-c) Variance is concentrated on L1, es27, e-Site tRNA, and the mRNA channel entrance at all resolutions. (d-f) Regular WMDs are dominated by noise at all but the lowest resolution. (h-j) Negative control shows no meaningful concentration of the variance due to severe missing wedge bias.

A rigid body docking of the full 80s mammalian ribosome in the non-rotated POST state from PDB-4UJE^64^ showed very clear agreement with the conformation of the 40s subunit, which combined with the co-factors observed suggest classes II and III are POST trans-locational ribosomes differing in retention of E site tRNA while class I is most similar to the “sampling” state. Classes IV and V both have eeF2 bound and differ in rotation of the 40s subunit of 5.9☐ and 5.0☐, respectively. Rigidly docking the 80s yeast structure of eeF2 from PDB-4UJO^65^ into classes IV and V show overall good agreement with their eeF2·sordarin·GDP position and our density. There are, however, noticeable differences, particularly in domain IV of eeF2 which is known to be dynamic and plays a key role in translocation^66,62^. We analyzed these differences qualitatively by comparing the rigidly docked model solved with Sordarin present (figure 4j-k) with the same model after running a short (1ns) Molecular Dynamics Flexible Fitting (MDFF) (figure 4h-i). The antibiotic Sordarin is highly specific for binding to fungal eeF2 and permits GTP hydrolysis, yet prevents conformational changes that result in subsequent release of eeF2 after translocation^67^. Although Sordarin is not present in the sample under study here, there is a pronounced difference in electron density between domains III-V of eeF2 in class V (figure 4i black arrow) that coincides with the Sordarin binding pocket. This density is not present in class IV which also exhibits a rotation of eeF2-domain IV (figure 4j).

The application of multi-scale clustering is illustrated in figure 4, where the arrows point to the absence or presence of es27 carrying different significance as revealed by the eigenimage they appear in. (These are the sum of the average and each eigenvector as determined by SVD.) Perhaps even more clear to see is the impact the 3D-CTF correction to WMDs has: at all length scales the variance maps shown in red in figure 4a-c clearly are concentrated on areas of true variance, in this case es27, L1, the e-Site and elements surrounding the mRNA tunnel entrance; regular WMDs only work at the lowest resolution while noise begins to dominate as higher resolution information is included, a necessary feature to fully resolve the classes shown; finally as a negative control the variance maps without any compensation are shown in figure 4g-i where the variance does not correspond to any relevant features.

## Discussion

With the rapid expansion of cryoEM resources available at major universities and with the development of shared use models like eBIC at Diamond Light Source, the ability to collect high quality cryoEM and cryoET data is now less limiting than the ability to effectively process the data. We have created a set of image processing routines incorporated into the program emClarity, which have demonstrated a much greater accuracy in alignment and image restoration compared to current state-of-the-art approaches using the same raw data sets which are publicly available. In addition to a pronounced enhancement in overall resolution, we have demonstrated a powerful approach for image classification in the presence of the “missing-wedge” effect by combining the correction for wedge differences with multi-scale clustering which helps to encode biologically relevant information for the clustering algorithms. The application of these advances to study samples free from biochemical restraints *in situ* shows promise, where, in addition to isolating known functional intermediates of translocation from a crude lysate, we also demonstrate the possibility of a more explorative approach. Having isolated classes IV and V of the mammalian ribosome *in situ* suggests both that Sordarin binding stabilizes an interaction between eeF2-domian III/V that exists on pathway in functional ribosomes and that nearby intermediates on the energy landscape, which are not observed in studies using the antibiotic, may be explored and observed by using this approach. In addition to isolating well-resolved class averages with minor populations, and finding nearby minima in the energy landscape, our approach also results in the production of accurate 3D-variance maps which will be beneficial to exploring macromolecular dynamics. By highlighting key regions of dynamic behavior, our approach should be useful for direct analysis and the design of complementary biophysical experiments. While these advances in classification are in the pre-processing and dimensionality reduction stage, future work to explore modern approaches in pattern recognition and machine learning, will likely establish another substantial improvement in the technique.

## Methods

Details for processing the RELION data set are available in the online tutorial on the emClarity wiki, which we will outline here. The tilt series were downloaded from EMPIAR, and a coarse alignment was obtained using the gold-fiducials in IMOD. Candidate particles were selected using template matching at an object pixel size of 13 Å (binning of 6) in emClarity with EMD-3228 low-pass filtered to 40 Å as a reference. After removing false positives using the 3D editing tools in emClarity, the defocus for each projection was then determined in emClarity and the CTF phase inversions were corrected on the projections. A six-parameter search at a binning of 4 for two iterations was carried out followed by an initial round of tomo-CPR where the local alignments were estimated. Since there are only 5-7 fiducials available for the coarse alignment, we returned to the template matching with the updated tilt-series alignment as this is the only full “global” search. Resuming the sub-tomogram workflow alternating 2 iterations of alignment followed by tomo-CPR at binning of 4, 3, then 2 each time re-interpolating the CTF corrected stacks following tomo-CPR. After the final round, we return to the raw data re-interpolating and then correcting the CTF on this stack. Following this we went through several iterations at full sampling to arrive at the final reconstruction. The processing of the mammalian data set was essentially the same.

Once the sub-tomograms were brought into alignment with a common reference frame, we ran classification using the full 3D-CTF to form the missing wedge compensated feature vectors, and at resolutions of 10, 16, 24 and 40Å for multi-scale PCA.

## Acknowledgments

We thank Dr. Joachim Frank and Dr. Wen Li for very helpful discussions. Mr. Doug Bevan for technical assistance with computer clusters, and Drs. Teresa Brosenitsch, Frances Joan-Alvarez, and J. Peter Rickgauer for reading the manuscript. We thank Dr. Xiaofeng Fu for testing the emClarity software. This work was supported by the National Institutes of Health (GM085043, GM082251) and the UK Wellcome Trust Investigator Award (206422/Z/17/Z).

## Author contributions

This study was conceived and designed by P.Z and B.A.H. B.A.H. developed and tested the code for emClarity. B.A.H. and P.Z. analyzed the results. B.A.H. and P.Z. wrote the paper.

## Additional information

### Accession codes

Cryo-EM structural data have been deposited in the EM Data Bank under accession codes EMD-8799 for the yeast 80s ribosome, EMD-8802, EMD-8803, EMD-8804, EMD-8805, and EMD-8806 for rabbit 80s ribosome classes I-V respectively.

## Supplemental Methods

### Refinement of tilt-series alignment

Tomo-CPR works by combining the strengths of the 3D-model-based and feature-tracking approaches, while also taking advantage of the robust alignment tools developed for gold-fiducial alignment available in the IMOD package. Starting with the tomogram (as in the 3D model approach), we additionally replace the density corresponding to our particles of interest at the proper orientation, with a copy of the high SNR sub-tomogram average and then re-project that synthetic tomogram using the IMOD program *tilt* supplemental figure 1a. This reprojection also includes any local alignments previously determined and allows us to create a reference tilt-series along with a model for each sub-tomogram position in the 2D-projection. Tiles around each projected sub-tomogram origin are masked out and multiplied by the CTF of the data projection at that point, explicitly considering the defocus gradient and the structural noise from each particle’s unique environment (supplemental figure 1b and 1c). These tiles are subsequently used as sub-tomogram “fiducials” to refine the positions via cross-correlation. These refined positions are then used as input to IMOD’s *tiltalign* as if they were derived from gold fiducials, allowing us to take advantage of local refinement and robust fitting, as described previously^40^. The global changes to the projection geometry are applied to the tilt-series, while the local refinements are taken into consideration when the tomograms are reconstructed on the fly by emClarity. In addition to the importance of considering neighboring particles (supplemental figure 2d), additional high-contrast features, like the edge of the carbon foil or other particulate matter are pointed out in figure 2e, where a thin strip of one of the tomo-CPR references, prior to tiling, is shown.

### Improved de focus determination _ for more accurate CTF correction

We devised a novel approach which extends this concept to resample *all* the CTF information by taking advantage of the fact that the discrete Fourier transform (dFFT) defines the spatial frequency (*q*) at any point in an image in a fashion that depends on the real space sampling rate (*S*) and image size (*N*) as in equation 3.

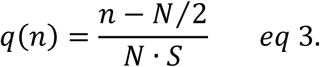

We first note that the spherical aberration term (*C_s_*) of the phase aberration equation (*χ_q_*) is roughly three orders of magnitude smaller than the defocus term, around 1nm^-1^ spatial frequency for defocus values around 3-4 μm.

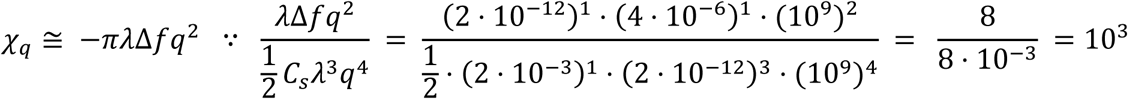

By considering this approximate expression, we suggest that the apparent defocus, i.e. where the CTF zeros end up in an image, can be effectively mapped from an image with defocus Δ*f*_0_ + ΔΔ*f* → Δ*f*_0_ by changing where the spatial frequency is sampled in the image of the transform, allowing all the tiles from a tilted projection to contribute coherently to the periodogram average and thereby permitting a per-tilt defocus determination. We find similar success by scaling with the right-hand term in equation 4 either the sampling rate in real space via interpolation, or equivalently scaling the image size via padding prior to computing the dFFT.

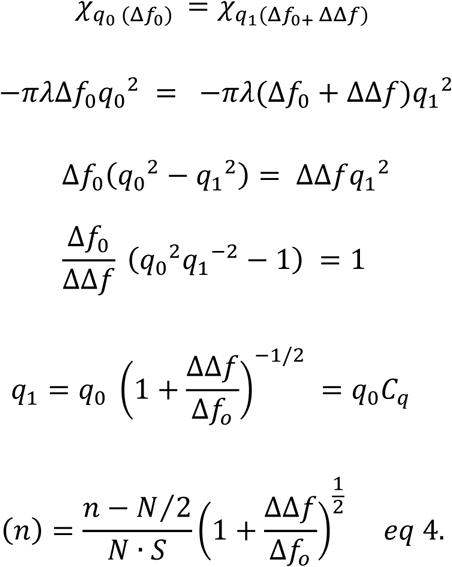

### Statistical optimization of the SNR in the final map

As it is necessarily a post-reconstruction filter, we start with the form of equation 8 in the original paper, and make modifications which explicitly consider anisotropy.

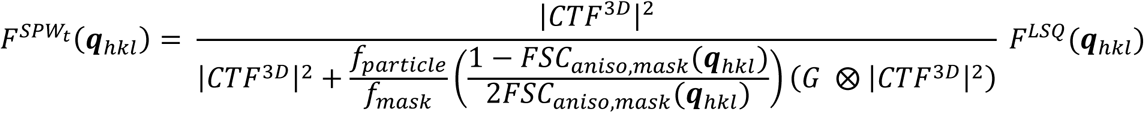

Where

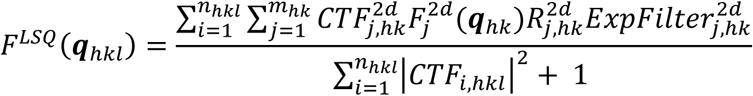

And

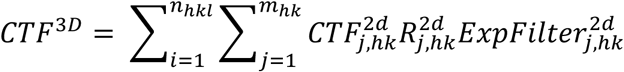

Here, the FSC normally calculated over spherical shells is further partitioned into shells limited to conical regions, which has been used previously to estimate resolution anisotropy^51^ and, in this way, the SSNR can be optimized for regions that may be less well sampled in reciprocal space. We also replace the average sampling over spherical shells, which is used to estimate the SSNR in the individual particles by one that is averaged over a local window using a simple Gaussian kernel for convolution. This is because under-sampled regions of the image may not necessarily have a low estimated SSNR which is a weakness in using the FSC for estimating this quantity.

### 3D-CTF compensated classification

For the special case where sub-tomograms are all oriented similarly, being adsorbed to a lipid monolayer for example, they may be averaged along the direction of their missing wedge and classified in 2D^52^; however, this is obviously of limited interest for most specimen. Another popular approach involves classifying the constrained-cross-correlation matrix^7,53^, however, this can be expensive to calculate and also disregards large amounts of information – discussed in detail in the paper on wedge masked differences (WMDs)^54^ on which we further expand and enhance.

The WMDs approach seeks to compensate the missing wedge by forming the difference between a given particle and its expected value. The expected value is estimated to be the global average distorted by the particle’s missing wedge. These are mean centered, normalized to a variance of one, and arranged into a 2D matrix followed by singular-value decomposition (SVD). The binary wedge used in this approach is only a first approximation, and we replace it with our full 3D-CTF. This correction allows the classification to include higher-resolution details than previously possible, which is a necessary but not sufficient condition to achieve the classification we report. We find that including the highest variance information from three to four discrete length scales at the same time is required to observe each class.

**Figure S1.**
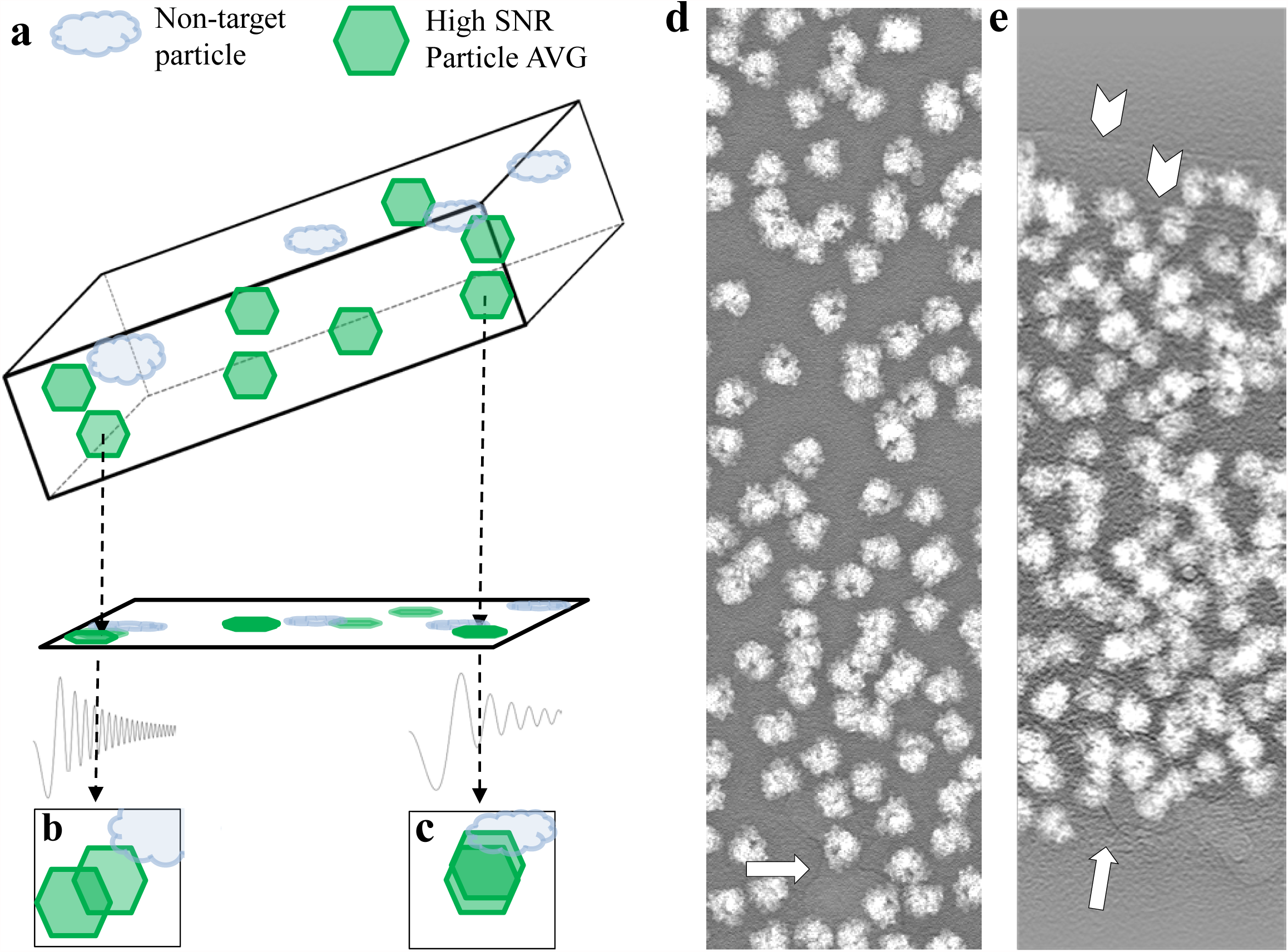
Tomo-CPR, constrained projection refinement. (a) Schematic overview for reference generation in the Tomo-CPR. The instances of structural noise resulting from the complex 3D environment of the sample are captured by projecting the full tomogram with the particle of interest replaced by the high SNR average. (b-c) Cartoon illustrating overlapping information in the projections arising from other particles, other components in the specimen, and variable defocus as a function of tilt. Examples of non-tilted (d) and tilted (e) projections used to generate references for the yeast 80s tomo-CPR. Tiles centered on the projection origin of each sub-tomogram are convolved with the CTF considering the individual sub-tomogram’s defocus, which varies with tilt angle and location in the tomogram. Particles are observed to overlap, even without tilting, while features due to contaminants (white arrows) and the carbon edge (white chevron) would prevent accurate determination of local particle drift using a simple projection of the average for crosscorrelation.

**Figure S2.**
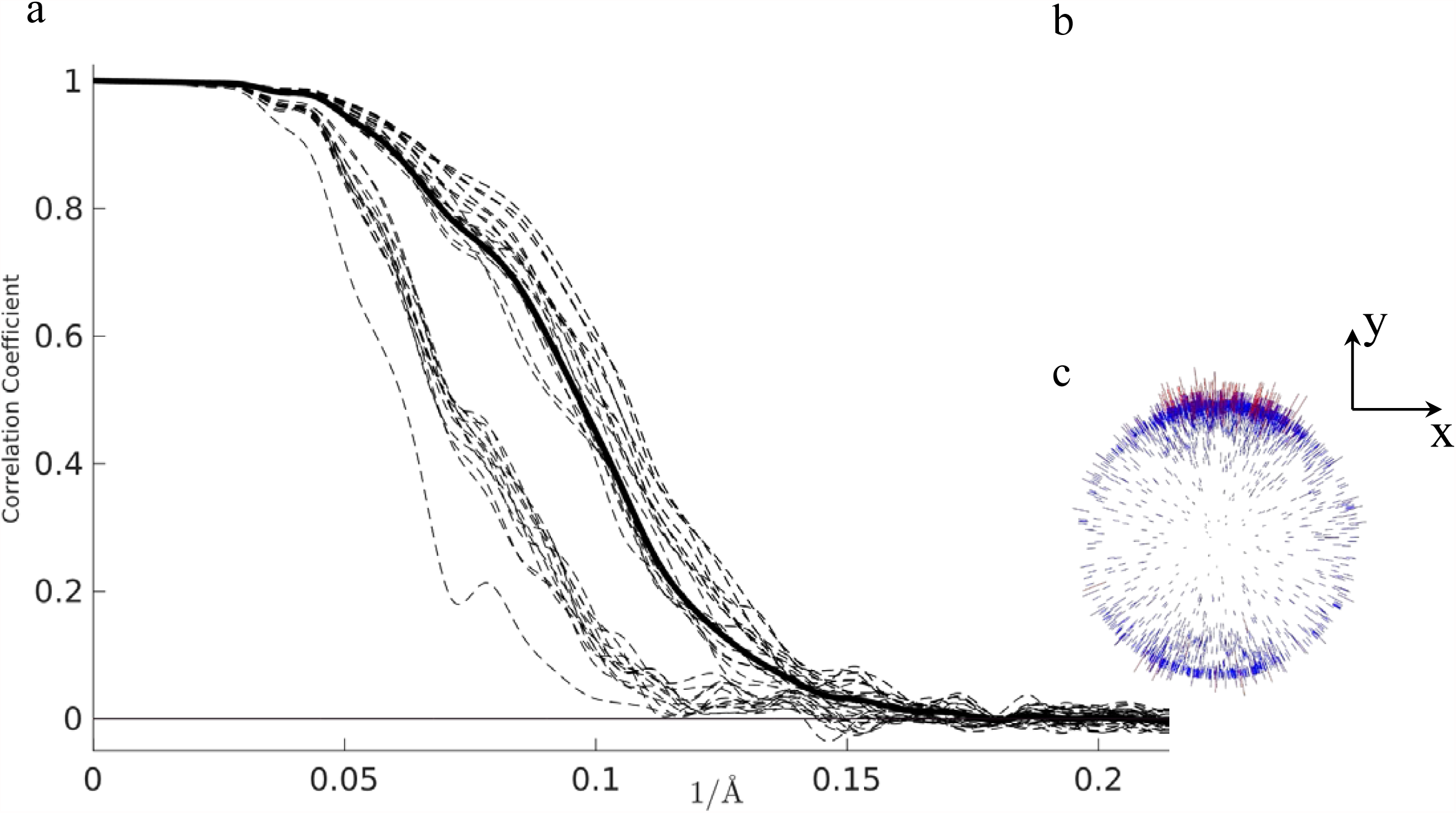
“gold-standard” FSC between half sets of yeast 80s ribosomes from EMPIAR-10045. (a) Plots show FSC calculated over 38 cones (dashed lines) with the overall average FSC in solid black. (b) Angular distribution plots for emClarity and (c) RELION. The microscope reference frame indicated with the Y-axis corresponding to the tilt-axis and in a right-handed convention. Given the particularly strong orientation preference, data along the Z-axis is of notably lower SSNR due to the missing-wedge effect and stands out from all the other FSC-cone curves.

## Footnotes

1 It is worth noting that the authors for the maps we use for comparison are also authors on the primary publications for their respective software packages, which helps to ensure the resolutions reported are likely optimal for the given data.

